# Role of polymetallic-nodule dependent fauna on carbon cycling in the eastern Clarion-Clipperton Fracture Zone (Pacific)

**DOI:** 10.1101/2022.06.21.496948

**Authors:** Tanja Stratmann

## Abstract

The abyssal seafloor in the Clarion-Clipperton Fracture Zone (CCZ) in the central Pacific is covered with large densities of polymetallic nodules, i.e., metal concretions containing iron, manganese, nickel, cobalt, and copper. These nodules are of economic interested and considered potential future resources for said metals, but they also host a variety of deep-sea fauna. In a recent study it was estimated that the removal of these nodules would lead to a loss of up to 18% of all taxa in the CCZ. Here, I assess the impact of removing these nodule-dependent taxa on carbon cycling at two sites (B4S03, B6S02) of the Belgian exploration license area in the eastern CCZ. For this purpose, I developed two highly-resolved carbon-based food web models with 72 (B6S02) to 77 (B4S03) food-web compartments consisting of different detritus pools, bacteria, metazoan meiobenthos, macrobenthic isopods, polychaetes and other macrobenthos, megabenthic cnidarians, crustaceans, poriferans, holothurians and other invertebrate megabenthos, and fish. These compartments were connected with 304 (B6S02) to 338 (B4S03) links which were reduced by 5–6% when nodule-dependent faunal compartments were removed. The models estimated the total system throughput *T*‥, i.e., the sum of all carbon flows in the food webs, in intact food webs as 1.24 mmol C m^−2^ d^−1^ and 1.20 mmol C m^−2^ d^−1^ at B4S03 and B6S02, respectively, whereupon 67.7% (B4S03) to 69.8% (B6S02) of *T*‥ flowed through the microbial loop. A removal of the nodule-dependent fauna did not affect this microbial loop, but reduced the scavenger loop by 54.6% (B6S02) to 84.1% (B4S03). Overall, nodule-dependent fauna is responsible for only a small fraction of total carbon cycling at the eastern CCZ. Therefore, when the effect of prospective deep-seabed mining on carbon cycling is investigated, its impact on benthic prokaryotes and the microbial loop should be addressed specifically.

## 1. Introduction

The abyss between 3,000 and 6,000 m water depth is the largest ecosystem on our planet (Gage and Tyler, 1991) and covers 85% of the seafloor (Harris et al., 2014). It consists of abyssal hills, abyssal mountains, and abyssal plains. The latter can have a difference in elevation between hilltops and valleys of 0 to 300 m and are often very flat with slopes of less than 0.05° (Cormier and Sloan, 2018). Abyssal plains are subdivided by mid-ocean ridges, like the East-Pacific Rise, deep-sea trenches, and island arcs, and interspersed with hills and mountains (Smith et al., 2008). In areas with well oxygenated bottom waters, high numbers of shark teeth, shells of plankton, or small rocks at the sediment surface, and sedimentation rates of less than 20 mm ky^−1^, abyssal plains are covered with polymetallic nodules, also known as manganese nodules (Hein et al., 2013; Hein and Koschinsky, 2014; Petersen et al., 2016). These nodules are concretions of iron oxide-hydroxide and manganese oxide (Hein and Koschinsky, 2014) and contain nickel, copper, cobalt, titanium, molybdenum, lithium, rare earth elements, and yttrium (Petersen et al., 2016). They form with growth rates of 1 to 10 mm million yr^−1^ (Petersen et al., 2016). Due to their high content of metals required for the transition from a fossil fuel-based economy to a renewable-energy based economy, polymetallic nodules are considered a potential future source of cobalt, copper, nickel, and rare earth elements (Hein et al., 2013).

Areas with polymetallic nodule densities of economic interest are the Central Indian Ocean Basin, the seabed around the Cook Islands, and the Clarion-Clipperton Fracture Zone (CCZ) (Kuhn et al., 2017). The CCZ is located south of Hawaii and west of Mexico in the central Pacific Ocean between 0°N, 160°W and 23.5°N, 115°W (International Seabed Authority, 2011). It has nodule densities between ∼1.5–3 kg m^−2^ in the south to ∼7.5 kg m^−2^ in the east (Washburn et al., 2021) and experiences a north-west (1.3 mg C_org_ m^−2^ d^−1^) to south-east (1.8 mg C_org_ m^−2^ d^−1^) gradient in particulate organic carbon flux (Vanreusel et al., 2016). These two environmental variables together with seafloor topography drive the different habitat classes in the CCZ (McQuaid et al., 2020).

Polymetallic nodules are not only interesting from an economic point of view, but also provide essential hard substrate for deep-sea benthos: Stalked sponges, such as *Hyalonema* (*Cyliconemaoida*) *ovuliferum* (Kersken et al., 2018), the encrusting sponge *Plenaster craigi* gen. nov. sp. nov. (Lim et al., 2017), and large xenophyophores (Gooday et al., 2017) grow on them. In fact, Veillette et al. (2007) identified 73 protozoan and 17 metazoan taxa attached to polymetallic nodules from the CCZ and Vanreusel et al. (2016) showed that even mobile benthos might benefit from the presence of nodules as their densities were higher in areas with nodules compared to nodule-free areas. To quantify the role of polymetallic nodules for trophic and non-trophic interactions in the CCZ, Stratmann et al. (2021) recently presented a highly-resolved interaction-web model: When the authors removed the nodules from the interaction web, they detected knock-down effects causing the loss of 17.9% of all taxa and 30.6% of all network links. Their detailed model estimated that 4% of all meiobenthos (i.e., benthos >32 µm), 50% of the macrobenthos (i.e., benthos >250 µm/ 500 µm), 45% of the invertebrate megabenthos (benthos >1 cm), and 0.5% of the fish were missing and the most impacted phyla were Bryozoa, Cnidaria, Platyhelminthes, and Porifera. Several of the taxa affected by nodule removal in the CCZ occur in the exploration license area of the Belgian company *Global Sea Mineral Resources* (GSR) in the eastern CCZ. Therefore, by evaluating which of the taxa that disappeared due to the knock-down effects are part of the present food-web model (short summary is presented in Table A.1), and by subsequently excluding these ones from the model, one can assess the impacts of polymetallic nodule removal on carbon cycling.

Short-term carbon cycling in the abyss is dominated by prokaryotes (Stratmann et al., 2018b; Sweetman et al., 2019), before the contribution of macrobenthos to phytodetritus cycling might increase after several days (Witte et al., 2003). In fact, carbon-based food-web models for deep-sea stations in the NE Atlantic, the NE Pacific, and the SE Pacific estimated that prokaryotes contribute >70% to total benthic respiration at said specific sites (de Jonge et al., 2020; Dunlop et al., 2016; Durden et al., 2017; van Oevelen et al., 2012). These carbon-based food web models are based on the principle of mass conservation and combine physiological parameters (e.g. assimilation and growth efficiency, secondary production, mortality, respiration), with site-specific flux constraints on carbon influx and loss, and biomasses of individual food-web compartments to calculate the carbon flows between compartments in the pre-defined topological food-web (Soetaert and van Oevelen, 2009; van Oevelen et al., 2010). They have been used previously to assess the potential recovery from a small-scale sediment disturbance in the SE Pacific (de Jonge et al., 2020; Stratmann et al., 2018a) and here, they will be applied to estimate changes in carbon flows depending on the presence and absence of nodule-dependent fauna at two sites of the GSR exploration license area in the CCZ.

I will assess (1) the potential small-scale variability in carbon cycling between two sites in the north-eastern CCZ that lay approximately 280 km apart, and (2) the potential reduction in total carbon cycling that the removal of nodule-dependent fauna from the abyssal food web might cause.

## 2. Materials & Methods

### 2.1 Study site

In the north-eastern CCZ, the GSR exploration license area stretches from 16°N, -128°E to 13°N, -122°E and includes three non-adjacent areas, the so-called B2, B4, and B6 areas (Fig. 1). The most western area B2 is located at ∼15°43’ N, -126°42’ E, the central area B4 is located at ∼14°6’ N, -125°52’ E, and the most eastern area B6 is located at ∼13°51’ N, -123°17’ E. Inside the B4 and B6 areas, two 10 × 20 km sampling sites, the B4S03 (14.112 N, -125.871 E) and B6S02 (13.894 N, -123.297 E) sites, were identified by (de Smet et al., 2017) based on differences in polymetallic nodule properties.

**Figure 1.**
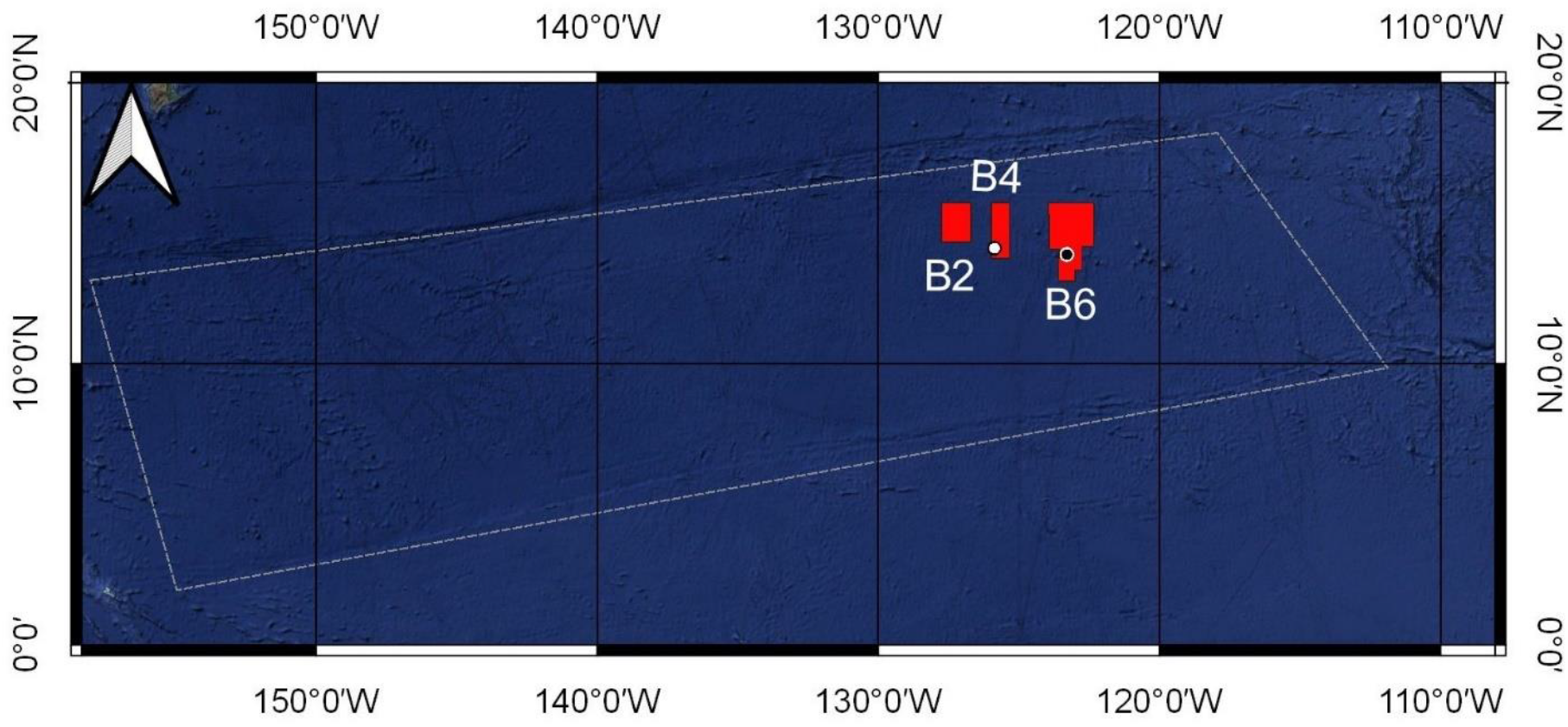
Map of the *Global Sea Mineral Resources* (GSR) exploration license areas inside the Clarion-Clipperton Fracture Zone (equatorial Pacific Ocean) with indications of the sites B4S03 (white dot) and B6S02 (black dot). The dashed polygon represents the CCZ and is based on the working definition of the CCZ by Glover et al. (2015). The coordinates of sub-part B6 are the revised coordinates as presented in International Seabed Authority (2019) and not the original coordinates included in the contract between GSR and the ISA.

Water depth at the two sites ranges between 4,470 and 4,560 m and the sediment is mainly muddy (mud content: 88.7–91.5%, sand content: 11.3–8.5%) with a median grain size of 16.5–18.5 µm (de Smet et al., 2017). Nodule density is 22.2±2.41 kg m^−2^ (±standard error) at B4S03 and 27.3±0.62 kg m^−2^ at B6S02 with a nodule surface coverage of 36.2±2.95% and 31.1±8.43%, respectively, and a nodule volume of 92.6±6.95 cm^3^ and 69.7±13.7 cm^3^ (de Smet et al., 2017).

### 2.2 Food-web structure

Food webs in the GSR exploration license area consisted of the abiotic compartments detritus (phytodetritus, semi-labile detritus, refractory detritus), dissolved organic carbon (DOC), and carrion, and the biotic compartments bacteria, metazoan meiobenthos (4 compartments), macrobenthos (40 compartments), invertebrate megabenthos (26 compartments), and fish (3 compartments). Metazoan meiobenthos and macrobenthic isopods were divided in feeding types based on peer-reviewed literature (Supplementary tables 1 and 2). The food-web compartments of all other macrobenthos, invertebrate megabenthos, and fish compartments resembled the highest taxonomic resolution possible, i.e., family level for macrobenthic polychaetes, (mainly) class level for arthropods, and phyla and class level for all other fauna.

**Table 1.**
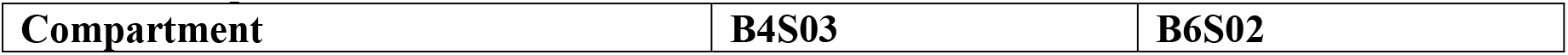
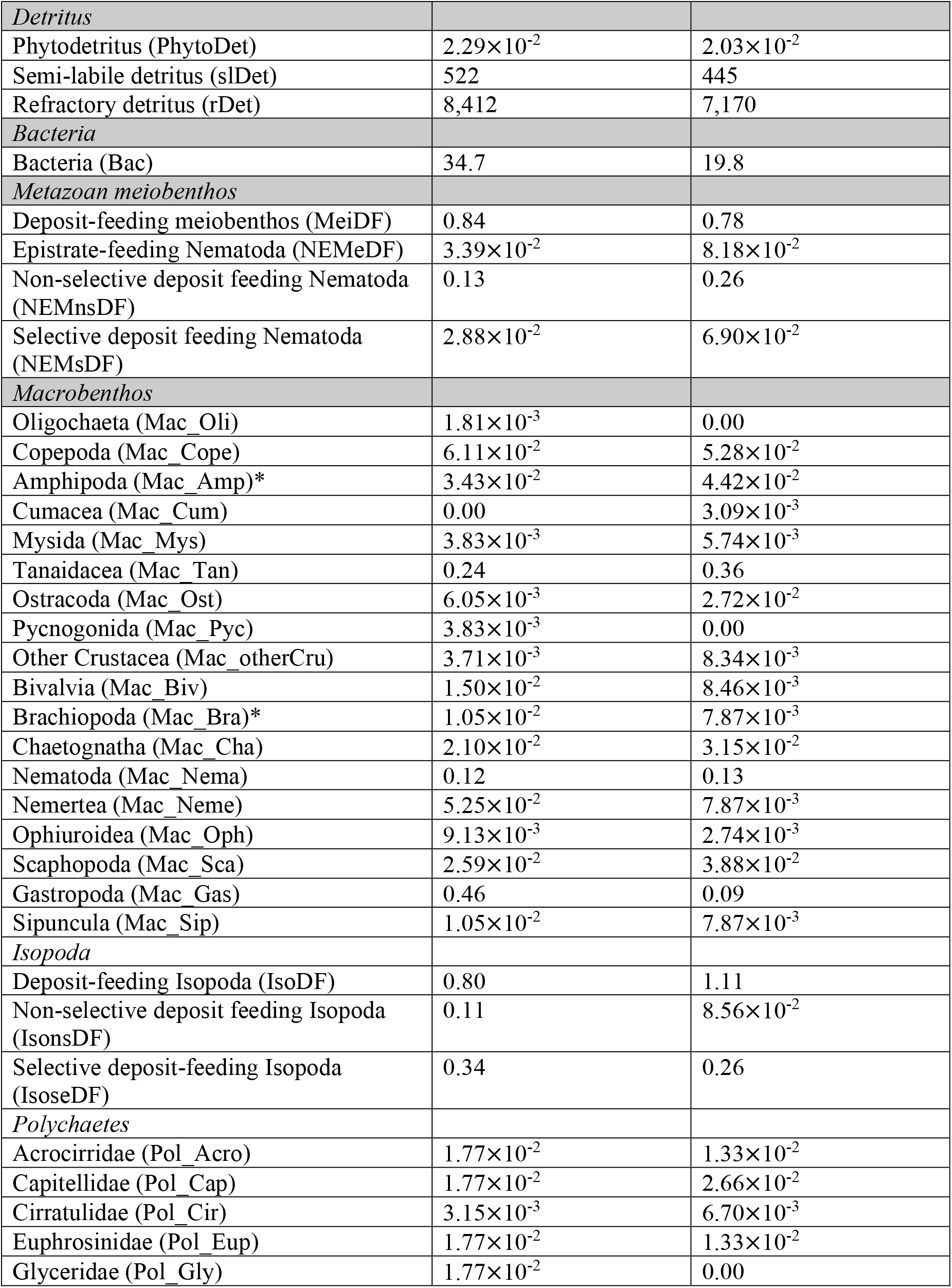
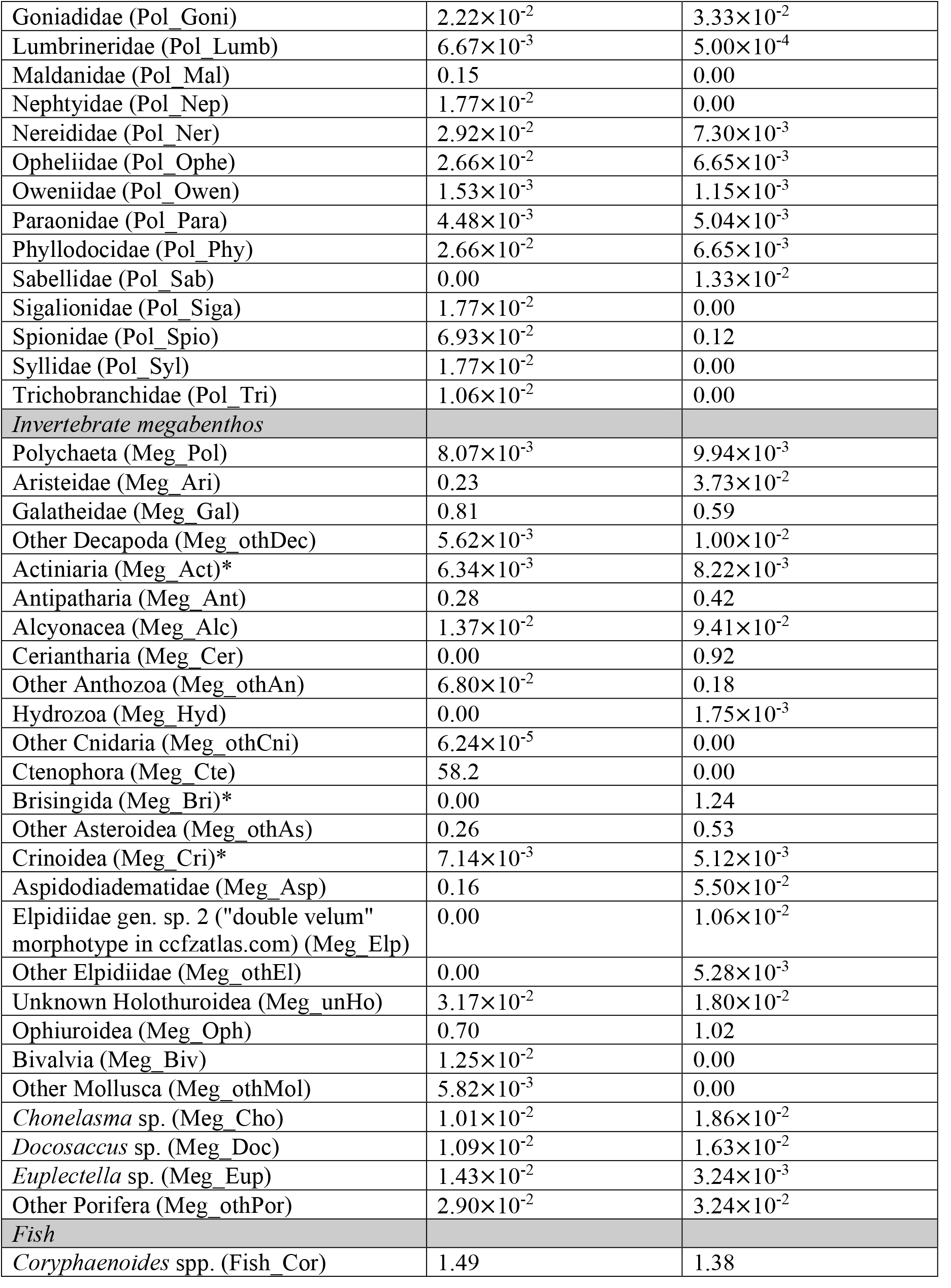
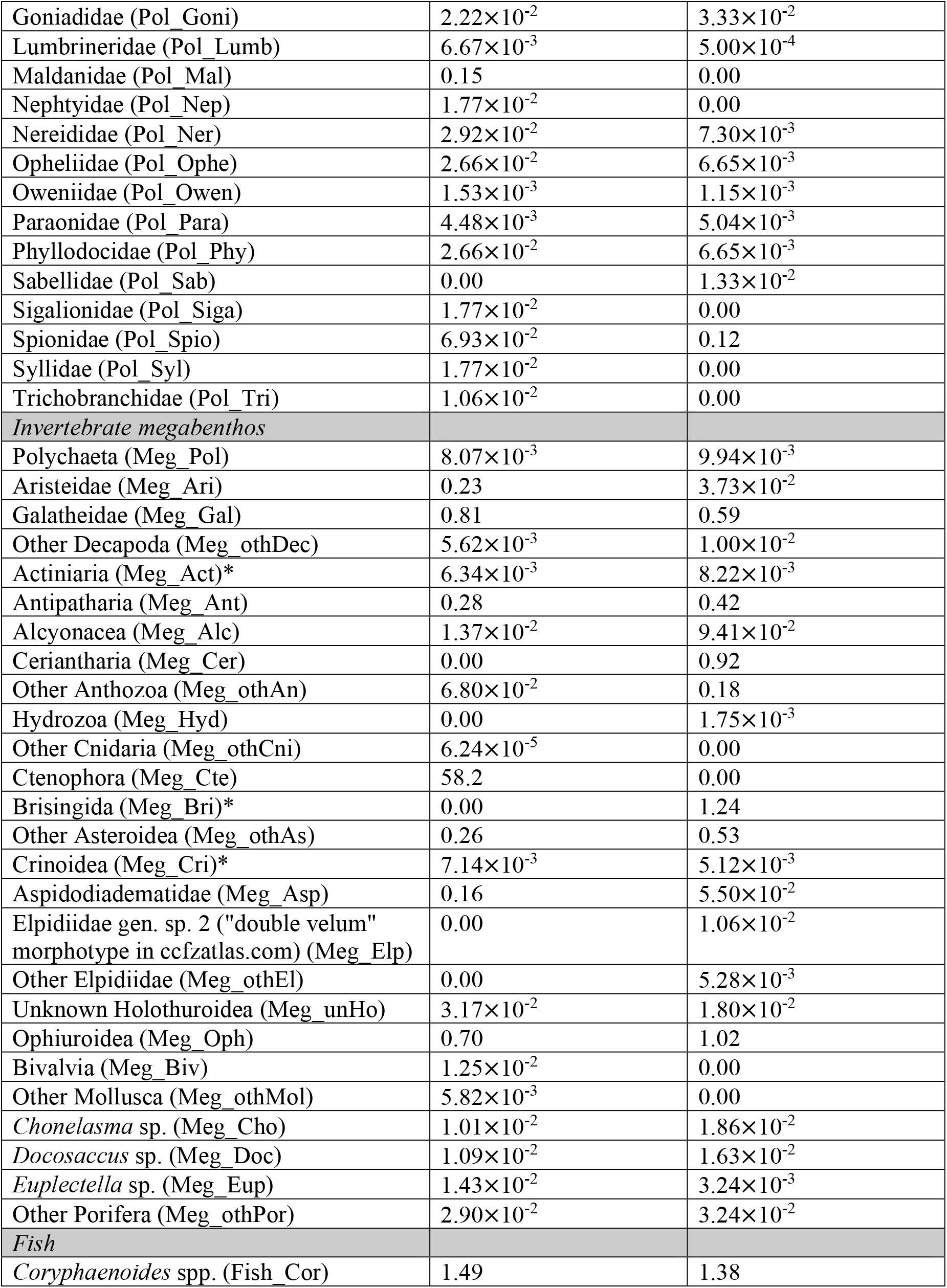
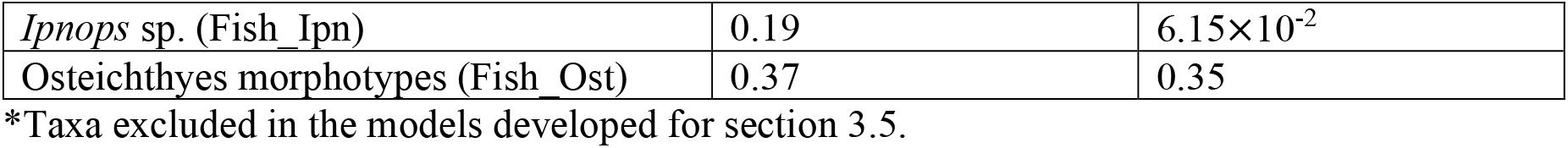
Carbon stocks (mmol C m^−2^) of all food-web compartments at sites B4S03 and B6S02 of the GSR exploration license area.

**Table 2.** Carbon flux data (mmol C m^−2^ d^−1^) from Volz et al. (2018) were implemented in the model as equalities (single value) or inequalities [min, max].

| Carbon flux | Value |
| --- | --- |
| Phydetritus deposition | [0, 0.118] |
| Semi-labile detritus deposition | [0, $7.945 \times 10^{-3}$] |
| Refractory detritus deposition | [$4.658 \times 10^{-4}$, ∞]* |
| Phydetritus degradation | lDet stock × [0, $5.479 \times 10^{-5}$] |
| Semi-labile detritus degradation | slDet stock × [$1.918 \times 10^{-8}$, ∞]* |
| Refractory detritus degradation | rDet stock × [$5.479 \times 10^{-10}$, ∞]* |
| Burial flux of refractory detritus | [0.013, ∞]* |
| Total C mineralization | [$7.75 \times 10^{-2}$, 0.115] |

\*Constraints adjusted in the fitting step.

Metazoan meiobenthos (>32 µm) included copepods and crustacean nauplii (Pape et al., 2017) that both deposit-feed on bacteria, phytodetritus, and semi-labile detritus (Supplementary table 1). Nematodes were classified as epistrate feeders, selective, and non-selective deposit feeders following Wieser (1953) based on the nematode genera present in the GSR exploration license area (Pape et al., 2017). They feed on bacteria, phytodetritus, and semi-labile detritus (Supplementary table 1).

Macrobenthos (>300 µm) contained annelids, arthropods, molluscs, brachiopods, chaetognaths, nematodes, nemerteans, echinoderms, and sipunculans (de Smet et al., 2017), that were identified to be carnivorous, (non-selective or subsurface) deposit feeders, filter- and suspension feeders, scavengers, and omnivores using peer-reviewed literature (Supplementary table 2). The polychaete feeding guild catalogue of Jumars et al. (2015) was inquired to classify the feeding types of polychaetes on family level. Isopods were grouped in the feeding types (selective, non-selective) deposit feeders based on the genera present in the GSR exploration license area (de Smet et al., 2017) using information on feeding type and food sources presented in Menzies (1962). Invertebrate megabenthos (>5 cm) comprised decapods, anthozoans and hydrozoans, poriferans, and echinoderms, i.e., crinoids, echinoids, holothurians, and ophiuroids (de Smet et al., 2021). These taxa were classified as deposit feeders, carnivorous suspension feeders, filter and suspension feeders, omnivores, and carnivores based on peer-reviewed literature (Supplementary table 3). The fish observed in the GSR exploration area were *Coryphaenoides* spp., *Ipnops* sp., and Osteichthyes morphotypes (de Smet et al., 2021) feeding on fish, crustaceans, polychaetes, and carrion (Supplementary table 3). A detailed list with taxon-specific feeding types and prey as well as food sources, respectively, is presented in Supplementary tables 1 to 3.

**Table 3.** Constraints of the physiological processes bacterial growth efficiency *BGE* (-), virus-induced bacterial mortality *VIBM* (-), bacteria carbon production *BCP* (mmol C m^−2^ d^−1^), assimilation efficiency *AE* (-), net growth efficiency *NGE* (-), secondary production *SP* (mmol C m^−2^ d^− 1^), mortality *M* (mmol C m^−2^ d^−1^), respiration *R* (mmol C m^−2^ d^−1^), and feeding preferences *FP* (-) implemented in the food-web models as equalities (single values) and inequalities ([minimum, maximum] values). In several cases, individual faunal constraints had to be adjusted in the fitting step. For these cases, the adjusted constraints are presented in Table A.2. References (Ref.): ^1^(Vonnahme et al., 2020), ^2^(de Jonge et al., 2020), ^3^(Danovaro et al., 2008), ^4^(Smith, 1978), ^5^(Drazen and Seibel, 2007), ^6^(Smith and Hessler, 1974), ^7^(Drazen and Yeh, 2012)

| Physiological process | Size class | Value | Ref. |
| --- | --- | --- | --- |
| *BGE* | Bacteria | [0.27, 0.90]* | 1, 2 |
| *VIBM* | Bacteria | [0.87, 0.91] | 3 |
| *BCP* | Bacteria | [0.34, 0.68] | 1,3 |
| *AE* | Metazoan meiobenthos | [0.18, 0.27] | 2 |
|  | Macrobenthos | [0.68, 0.89] | 2 |
|  | Invertebrate megabenthos | [0.40, 0.75] | 2 |
|  | Fish | [0.84, 0.87] | 2 |
| *NGE* | Metazoan meiobenthos | [0.10, 0.96] | 2 |
|  | Macrobenthos | [0.57, 0.68] | 2 |
|  | Invertebrate megabenthos | [0.23, 0.61] | 2 |
|  | Fish | [0.37, 0.71] | 2 |
| *SP* | Metazoan meiobenthos | $[2.00 \times 10^{-2}, 0.12] \times C \text{ stock}$ | 2 |
| | Macrobenthos | $[2.57 \times 10^{-3}, 1.67 \times 10^{-2}] \times C \text{ stock}$ | 2 |
| | Invertebrate megabenthos | $[3.18 \times 10^{-4}, 1.47 \times 10^{-3}] \times C \text{ stock}$ | 2 |
| | Fish | $[0, 6.30 \times 10^{-4}] \times C \text{ stock}$ | 2 |
| *M* | Metazoan meiobenthos | $[0, 0.12] \times C \text{ stock}$ | 2 |
| | Macrobenthos | $[0, 1.67 \times 10^{-2}] \times C \text{ stock}$ | 2 |
| | Invertebrate megabenthos | $[0, 1.47 \times 10^{-3}] \times C \text{ stock}$ | 2 |
| | Fish | $[0, 6.30 \times 10^{-4}] \times C \text{ stock}$ | 2 |
| *R* | Metazoan meiobenthos | $[7.00 \times 10^{-3}, 0.15] \times C \text{ stock}$ | 2 |
| | Macrobenthos | $[3.57 \times 10^{-5}, 4.21 \times 10^{-2}] \times C \text{ stock}$ | 2 |
| | Invertebrate megabenthos | $[9.32 \times 10^{-8}, 1.26 \times 10^{-3}] \times C \text{ stock}$ | 2 |
| | *Coryphaenoides* spp., Osteichthyes morphotypes | $[3.12 \times 10^{-4}, 4.84 \times 10^{-4}] \times C \text{ stock}$ | 4-7 |
| | *Ipnops* sp. | $[1.79 \times 10^{-4}, 8.54 \times 10^{-4}] \times C \text{ stock}$ | 2 |
| *FP* |  | [0.75, 1.00] | 2 |

\*Constraint adjusted in the fitting step.

### 2.3 Food-web links

Links of carbon transfer between the food-web compartments were implemented as presented in Fig. 2.

**Figure 2.**
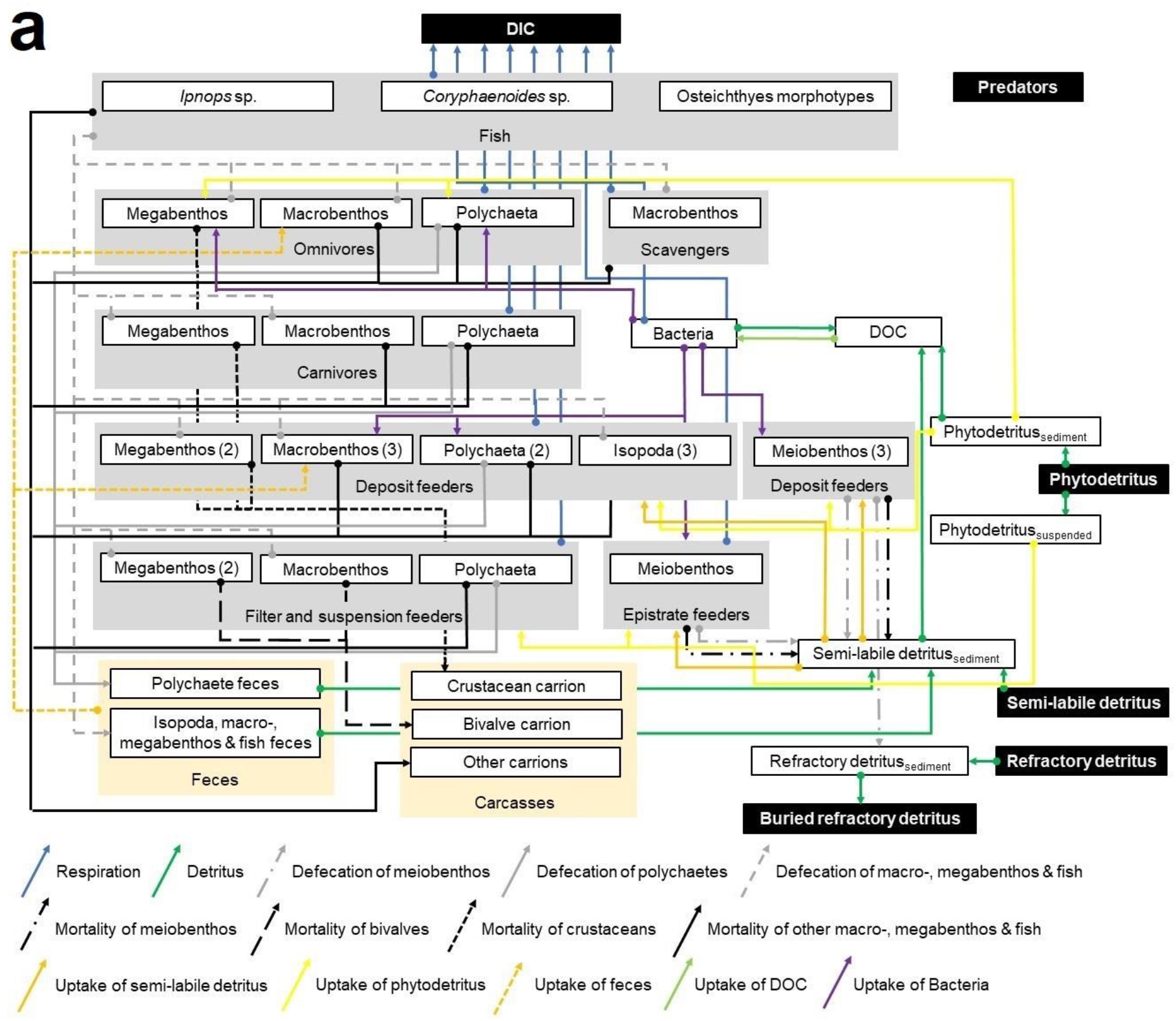
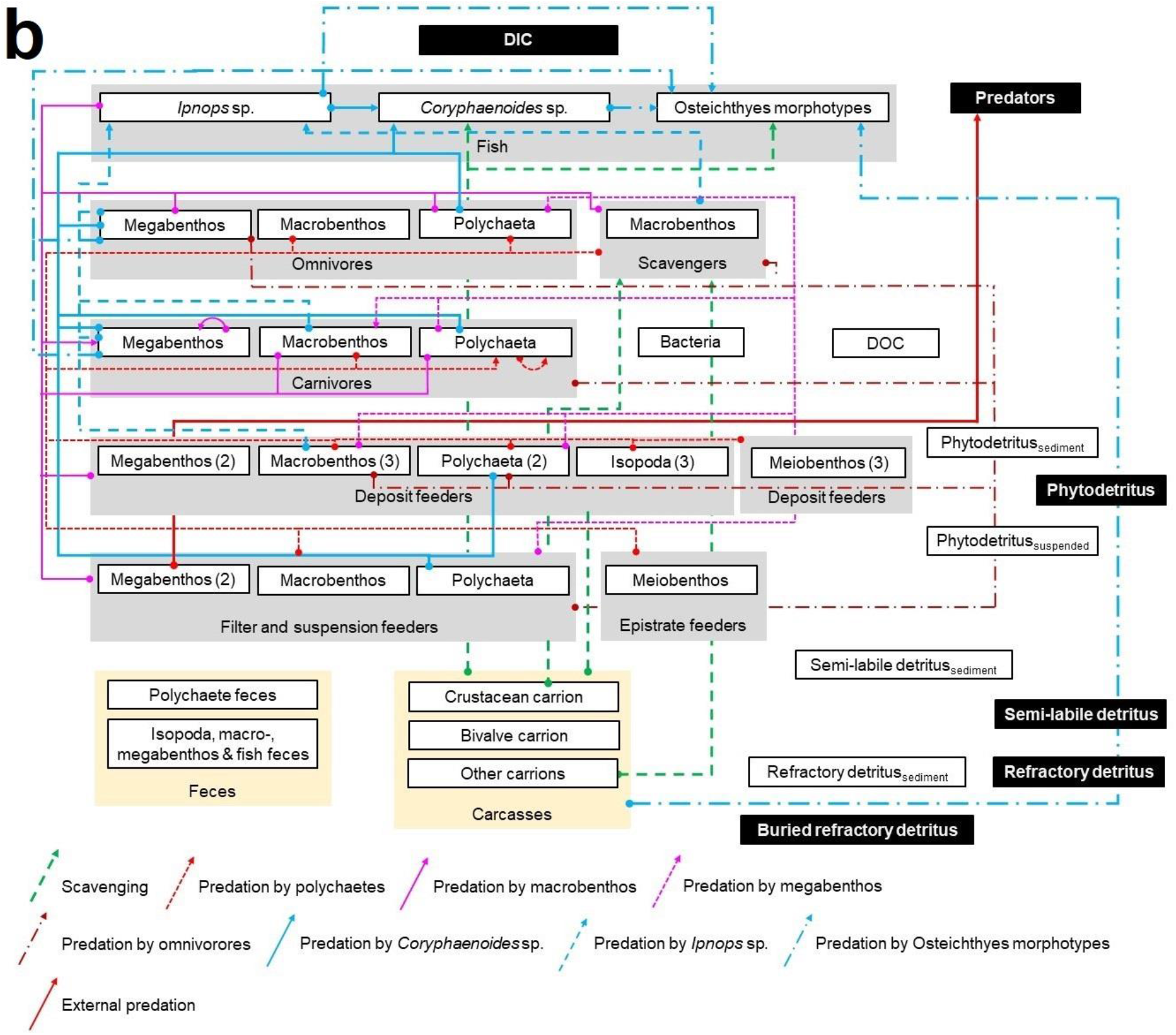
Presentation of the topological food web implemented in the linear inverse models for the GSR exploration license area. White boxes represent all compartments of the food web inside the model, in contrast to black boxes that show compartments outside the model that were not specifically modelled. Gray boxes display the feeding types epistrate feeders, filter- *and* suspension feeders, deposit feeders, carnivores, omnivores, scavengers, and fish. Yellow boxes represent feces and carcasses. White boxes enclosed by gray boxes show the size classes meiobenthos, macrobenthic isopods and polychaetes, other macrobenthos, invertebrate megabenthos, and the three fish taxa *Ipnops* sp., *Coryphaenoides* spp., and Osteichthyes morphotypes. Numbers in brackets indicate how many different kinds of a specific feeding type exist for a particular size class. Arrows show the flow of carbon from the carbon source (arrow ending with a big dot) to the carbon sink (head of arrow). The sedimentary phytodetritus pool receives carbon input via degrading phytoplankton in marine snow, the semi-labile and refractory detritus pools are maintained by semi-labile and refractory marine snow particles. Furthermore, fecal pellets, that are not taken up by specialized coprophagus feeders (i.e., organisms feeding on feces), contribute to the semi-labile and refractory detritus pool. Also dying metazoan meiobenthos becomes part of the semi-labile detritus pool. Dead macrobenthos, invertebrate megabenthos, and fish, in contrast, add to the carrion pool. All three detritus pools hydrolyze to DOC that is taken up by bacteria. Bacteria convert this DOC to dissolved inorganic carbon (DIC) via respiration and they contribute to the DOC pool when they burst due to virus-induced bacterial lysis. Carbon is lost from the modeled system via burial of refractory detritus, respiration of all fauna and bacteria, and by external predators that predate upon invertebrate megabenthos.

Figure panel (a) shows respiration in the form of dissolved inorganic carbon (DIC), fluxes of detritus, DOC, and feces, mortality of fauna, and uptake of Bacteria, detritus, and DOC. Figure panel (b) presents scavenging and predation.

Deposit feeders graze upon bacteria, semi-labile detritus, and phytodetritus. Suspension and filter feeders filter, depending on the taxon, marine snow, crustaceans of macrobenthic size and fish out of the water column. Porifera, i.e., *Chonelasma* sp., *Docosaccus* sp., *Euplectella* sp., and other unidentified poriferans, furthermore, take up marine snow and DOC. Carnivores predate upon their specialized prey reported in Supplementary tables 2 and 3, except for the polychaetes of the family Sigalionidae, for which no prey types were published. Hence, this compartment was defined to feed on prey of the same size class. The diet composition of the individual fish taxa was constrained based on the contribution of each prey taxon to the total prey carbon stock (Table A.3); the contribution of carrion to the fish diet, however, was not further constricted.

### 2.4 Data sources

#### 2.4.1 Carbon stocks of food-web compartments

Data on carbon stocks of the various food-web compartments were extracted from the literature and standardized to mmol C m^−2^ as described below.

The phytodetritus carbon stock in the upper 5 cm of sediment was calculated by converting the chlorophyll content (3.80×10^−4^ µg g dry mass sediment^−1^, (Pasotti et al., 2021)) measured by high performance liquid chromatography to carbon units using a carbon:chlorophyll a-ratio of 40 (de Jonge, 1980) and a porosity of 0.858 (B4S03) and of 0.874 (B6S02) (de Smet et al., 2017) assuming a dry sediment density of 2.55 g cm^−3^. Unfortunately, no chlorophyll a concentration data were available for B6S02, so that the concentration from B4S03 was also used for B6S02. No data on semi-labile detritus, here defined as the so-called biopolymeric carbon (Fabiano et al., 1995), i.e., the sum of proteins, carbohydrates, and lipids, existed for the GSR exploration license area or any other license area in the CCZ. Therefore, the sum of the semi-labile and refractory detritus carbon stock was calculated by subtracting the phytodetritus pool from the total organic carbon (TOC) stock based on TOC data from de Smet et al. (2017). Subsequently, the relation of semi-labile to refractory detritus of 1:17.3 from the Peru Basin (SE Pacific) (de Jonge et al., 2020) was used to calculate both carbon stocks.

Bacterial carbon stock in the upper 5 cm of sediment was measured by Pape et al. (2017) as polar lipid concentrations following the Bligh and Dyer method (Bligh and Dyer, 1959; Boschker et al., 1999) and subsequently converted to bacterial biomass by the authors using conversion factors from Middelburg et al. (2000) and Brinch-Iversen and King (1990).

Metazoan meiobenthos carbon stock in the upper 5 cm of sediment was calculated by multiplying meiobenthos densities determined in multicorer samples (inner core diameter: 10 cm; n = 3) by Pape et al. (2017) with taxon-specific individual biomass data from Supplementary table 1. Macrobenthos carbon stock in the upper 10 cm of sediment was estimated by multiplying macrobenthos density data taken from box corer deployments (surface: 0.25 m^2^, sediment depth: 60 cm; n = 4 at B4S03, n = 3 at B6S02) by de Smet et al. (2017) with taxon-specific individual biomass data from Supplementary table 2. Invertebrate megabenthos and fish density data were taken from an autonomous underwater vehicle megafauna survey of the seafloor in 6 m altitude (de Smet et al., 2021) and multiplied with taxon-specific individual biomasses from Supplementary table 3 to estimate invertebrate megabenthos and fish carbon stocks. A complete list with carbon stocks of all food-web compartments is presented in Table 1.

#### 2.4.2 Site-specific flux constraints

Site-specific flux constraints were estimated by Volz et al. (2018) in a numerical diagenetic model for the GSR exploration license area and were implemented in the food-web models as presented in Table 2.

Phytodetritus deposition corresponds to labile detritus deposition in Volz et al. (2018) and semilabile and refractory detritus deposition rates are the influx of semi-labile and refractory marine snow particles. Phytodetritus and semi-labile detritus degradation processes describe the reduction of these detritus pools by dissolution of the detritus to DOC and their uptake by fauna. Refractory detritus is degraded via dissolution of this pool to DOC and it is buried in the sediment via the burial flux. Total C mineralization relates to aerobic respiration that is responsible for 90% of the organic matter consumption at the seafloor in the CCZ (Volz et al., 2018).

#### 2.4.3 Physiological constraints

Constraints of physiological processes that were included in the model are presented in Table 3.

Bacterial growth efficiency *BGE* is defined as

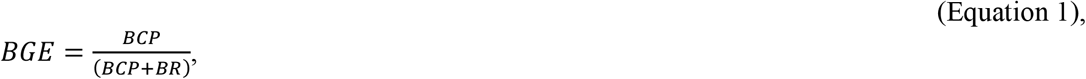

with *BCP* being bacterial carbon production (mmol C m^−2^ d^−1^) and *BR* being bacterial respiration (mmol C m^−2^ d^−1^).

Assimilation efficiency *AE* is defined as

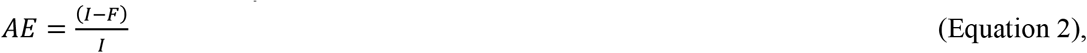

where *I* is food ingested and *F* relates to the feces produced (Crisp, 1971).

Net growth efficiency *NGE* is defined as

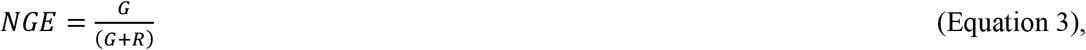

where *G* is the growth and *R* is the respiration (Clausen and Riisgård, 1996). The secondary production *SP* (mmol C m^−2^ d^−1^) is calculated as

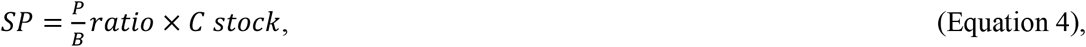

with 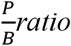 being the production/ biomass ratio (d^−1^).

The mortality *M* (mmol C m^−2^ d^−1^) always ranges from 0 to the maximum secondary production. Respiration *R* is calculated similar to secondary production as:

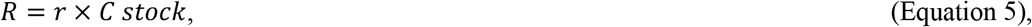

where *r* corresponds to the biomass-specific faunal respiration (d^−1^).

Feeding preferences *FP* of mixed deposit feeders *and* carnivores and omnivores indicate the contribution of predation to their respective diets.

### 2.4 Linear inverse model development and network indices

Carbon-based linear inverse food-web models were developed for steady state conditions based on the topological food web presented in Fig. 2. Said food-web models consist of linear functions in the form of equality (equation 6) and inequality matrix equations (equation 7) (van Oevelen et al., 2010):

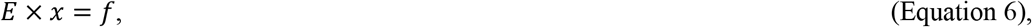

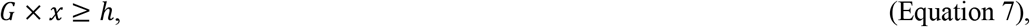

with vector *x* including the unknown fluxes. The vectors *f* and *h* include empirical equality and inequality data, and the coefficients in the matrices *E* and *G* describe combinations of unknown fluxes that have to fulfill the requirements defined in the vectors *f* and *h*.

Models for the two sites consisted of 654 (B6S02)–829 (B4S03) flows, 74 (B6S02)–79 (B4S03) mass balances (i.e., food-web compartments), 0 data equalities, and 340 (B6S02)–371 (B4S03) data inequalities. This meant that the models were mathematically under-determined (74–79 equalities vs. 654–829 unknown flows). In this case, either a single ‘best’ solution can be calculated based on the principle of parsimony or simplicity, or the uncertainty of each flow can be quantified using the likelihood approach (van Oevelen et al., 2010). Generally, the likelihood approach is recommended (van Oevelen et al., 2010), but this can be extremely computational demanding when the food-web models are exceptionally complex (see e.g. de Smet et al. (2016)). In fact, a test run on the bioinformatics server of the NIOZ Royal Netherlands Institute for Sea Research (The Netherlands) with 2,000 solutions calculated in one session required more than 1.5 months of calculation. As a result, the author decided to calculate the parsimonious solution of each food-web model and estimate ranges with the R package *LIM* v.1.4.6 (van Oevelen et al., 2010) in *R* v.1.3.1073 (R-Core Team, 2017).

The network indices “number of links” *L*_*total*_, “link density” *LD*, “connectance” *C*, “total system throughput” *T*‥ which is the sum of all carbon flows in the food web, Finn’s Cycling Index *FCI*, and the trophic level of each compartment were calculated using the R package *NetIndices* v.1.4.4 (Kones et al., 2009).

The trophic levels of the carrion pools were calculated as described in de Jonge et al. (2020) as:

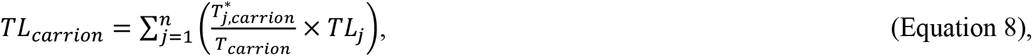

where *n* corresponds to the number of internal food-web compartments, *j* is all food-web compartments, *T\** is the flow matrix without external flows, and *T*_*carrion*_ is the total carbon inflow to the carrion pool without external sources.

## 3. Results

### 3.1 Benthic biomass

Total carbon stocks were 9,036 mmol C m^−2^ at site B4S03 and 7,646 mmol C m^−2^ at B6S02 (Table 1). Most of these carbon stocks consisted of semi-labile (B4S03: 5.78%, B6S02: 5.82%) and refractory detritus (B4S03: 93.1%, B6S02: 93.8%). Phytodetritus, bacteria, and metazoan fauna accounted for 2.53×10^−4^% (B4S03)–2.66×10^−4^% (B6S02), 0.38% (B6S02)–0.26% (B4S03), and 0.74% (B4S03)–0.14% (B6S02) of the total carbon stock.

The meiobenthic carbon stock contained 0.19 mmol C m^−2^ and 0.41 mmol C m^−2^ Nematoda at B4S03 and B6S02, respectively, and 0.84 mmol C m^−2^ and 0.78 mmol C m^−2^ other meiobenthos (i.e., copepods, nauplii) (Fig. 3). The carbon stock of macrobenthic Isopoda was 1.25 mmol C m^− 2^ and 1.46 mmol C m^−2^ at the two different sites and macrobenthic polychaetes had a carbon stock of 0.47 mmol C m^−2^ and 0.25 mmol C m^−2^ at B4S03 and B6S02, respectively (Fig. 3). The other macrobenthos carbon stock included mainly macrobenthic Nematoda, Tanaidacea, and Gastropoda, that contributed between 22.9% (B6S02) and 29.3% (B4S03) to the total carbon stock of this size class. The megabenthic carbon stock accounted for 60.9 mmol C m^−2^ at the B4S03 site and 5.23 mmol C m^−2^ at the B6S02 site (Fig. 3), whereupon at the first site 58.2 mmol C m^−2^ consisted of Ctenophora. Without this phylum, carbon stocks were 5.71 mmol C m^−2^ and 5.23 mmol C m^−2^ at the respective sites. Fish had a carbon stock of 2.05 mmol C m^−2^ and 1.79 mmol C m^−2^ (Fig. 3).

**Figure 3.**
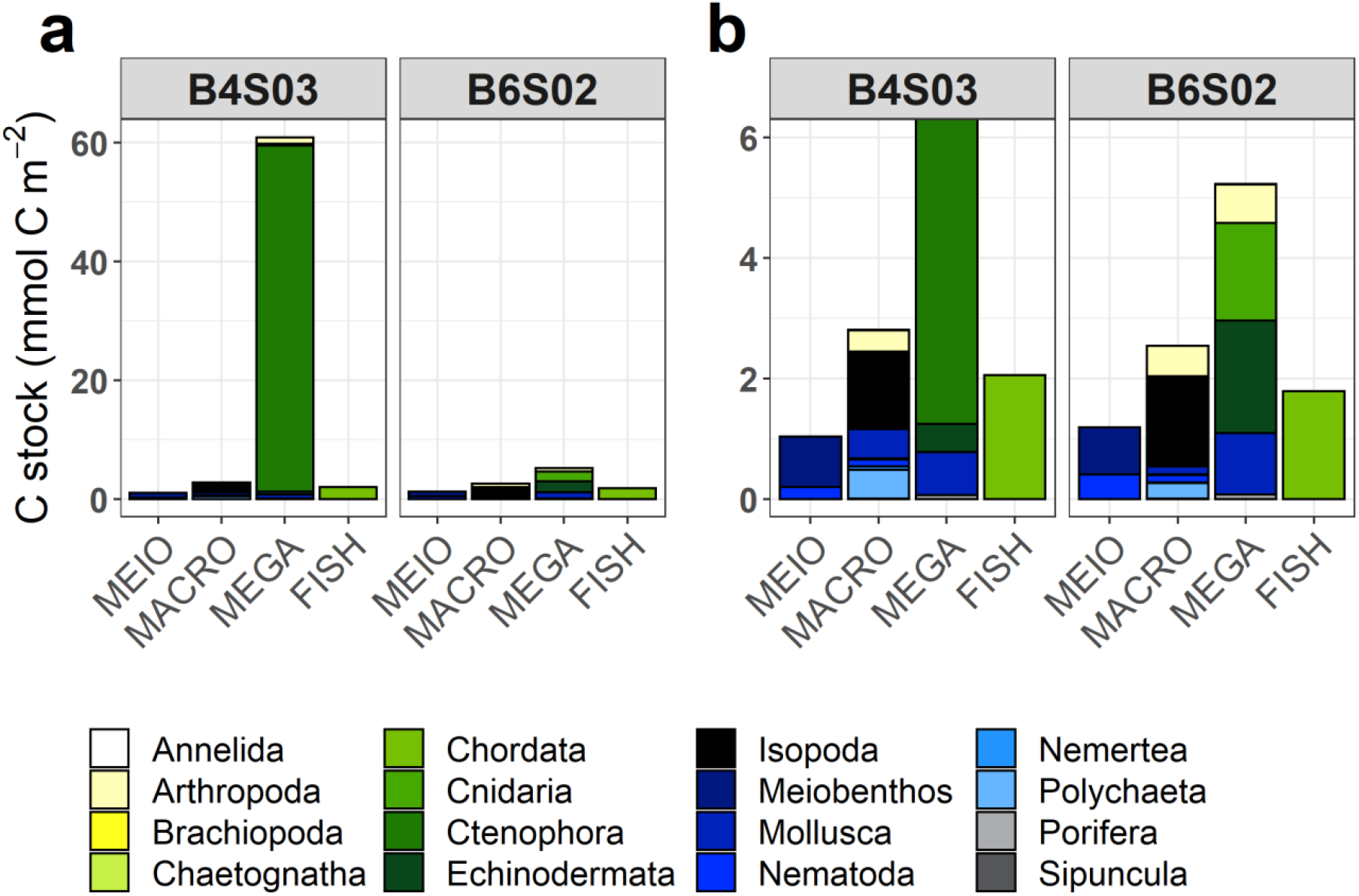
Mean carbon stocks (mmol C m^−2^) of the faunal food-web compartments at sites B4S03 (left columns) and B6S02 (right columns) in the GSR exploration license area. Panel (a) shows carbon stocks of all phyla, whereas for plotting purposes panel (b) shows only carbon stocks up to 6 mmol C m^−2^.

### 3.2 Food-web structure and trophic levels

The food webs consisted of 77 and 72 compartments at sites B4S03 and B6S02 that were connected with 338 and 304 links, respectively (Fig. 4, Table 4), resulting in a link density of 4.39 at B4S03 and of 4.22 at B6S02. The connectance was 4.51×10^−2^ at B4S03 and 4.62×10^−2^ at B6S02.

**Table 4.** Network indices calculated for the sites B4S03 and B6S02 when all fauna are present (+nodule) and when fauna predicted to be lost when polymetallic nodules are removed are excluded (-nodule). Relative changes (%) in network indices due to faunal loss are reported for each site. Abbreviations are: *n* = number of food-web compartments, *L*_*total*_ = number of links, *LD* = link density, *C* = connectance, *T*‥ = total system throughput, *FCI* = Finn’s Cycling Index.

| Network indices | B4S03 site |  |  | B6S02 site |  |  |
| --- | --- | --- | --- | --- | --- | --- |
|  | (+nodule) | (-nodule) | % relative change | (+nodule) | (-nodule) | % relative change |
| $n$ | 77 | 73 | -5.19 | 72 | 67 | -6.94 |
| $L_{total}$ | 338 | 308 | -8.88 | 304 | 290 | -4.61 |
| $LD$ | 4.39 | 4.22 | -3.88 | 4.22 | 4.33 | +2.51 |
| $C$ | $4.51 \times 10^{-2}$ | $4.53 \times 10^{-2}$ | +0.37 | $4.62 \times 10^{-2}$ | $5.13 \times 10^{-2}$ | +11.2 |
| $T_{..}$ | 1.24 | 1.24 | $-2.85 \times 10^{-2}$ | 1.20 | 1.20 | $+6.93 \times 10^{-2}$ |
| $FCI$ | 0.51 | 0.51 | -0.34 | 0.53 | 0.53 | +0.18 |

**Figure 4.**
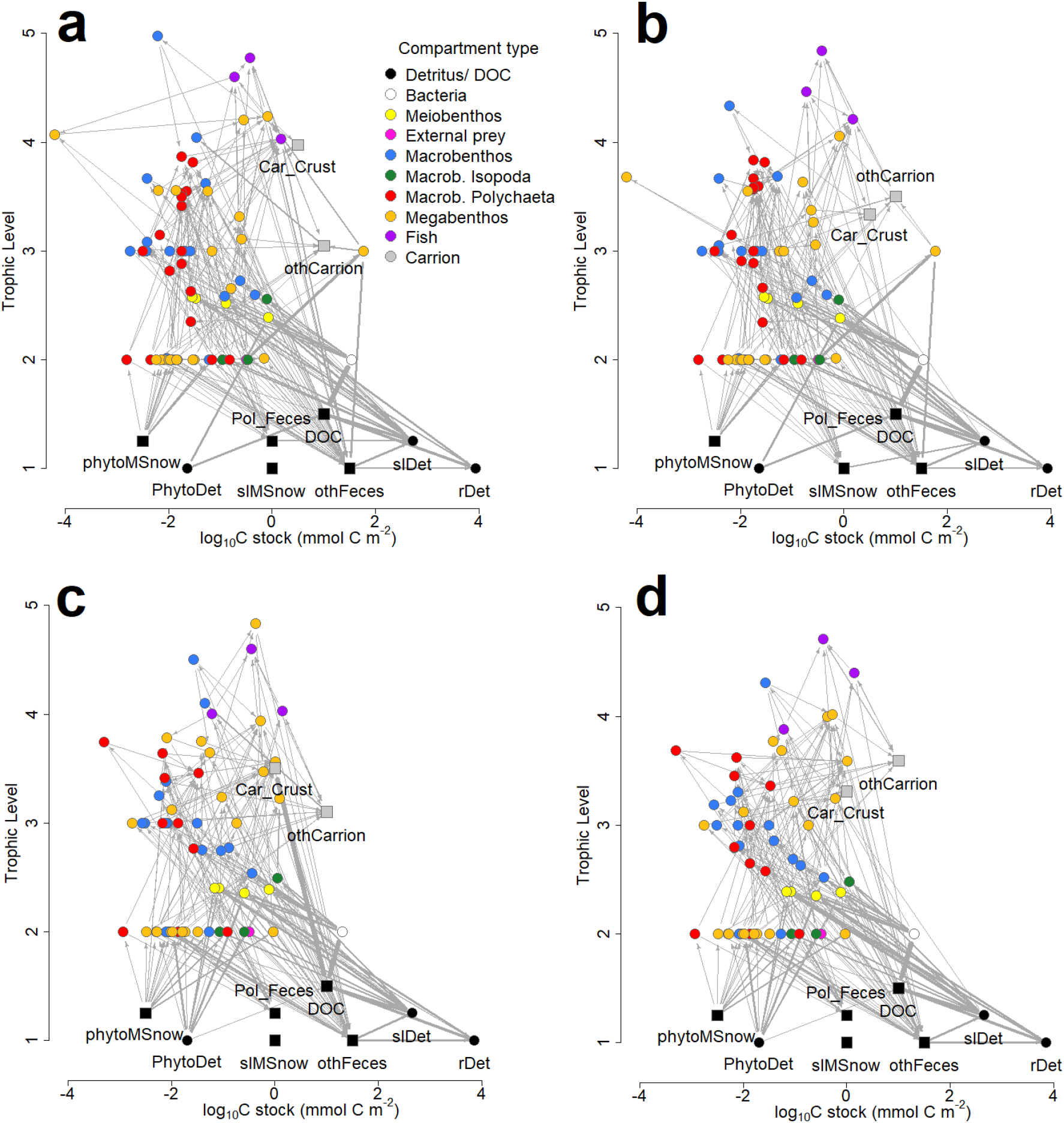
Structure of food-web models developed for the B4S03 site (a) in the presence and (b) absence of polymetallic nodules and for the B6S02 site (c) in the presence and (d) absence of nodules in the GSR exploration license area. Round nodes show compartments whose biomass (mmol C) is known, whereas square nodes indicate compartments to which the author assigned values for plotting purposes. Trophic levels 1.25 and 1.5 were assigned to square nodes for plotting purposes only; their true trophic levels are 1. The thickness of the arrows corresponds to the magnitude of carbon flow (mmol C m^−2^ d^−1^) between two food-web compartments after double square root-transformation. Note, that neither import nor export fluxes were plotted and that the x-axis is a log_10_ scale. Abbreviations are: Car_Crust = crustacean carrion, DOC = dissolved organic carbon, othCarrion = other carrion, othFeces = isopod, macro-, invertebrate megabenthic and fish feces, PhytoDet = phytodetritus, phytoMSnow = phytodetritus-based marine snow, Pol_Feces = polychaete feces, rDet = refractory detritus, slDet = semi-labile detritus.

Maximum trophic level was estimated as 4.97 (macrobenthic Ostracoda) at site B4S03 and as 4.83 (megabenthic Anthozoa) at B6S02. Mean modelled trophic level of metazoan fauna ranged from 2.84±0.10 (±standard error; B6S02) to 2.87±0.10 (B4S03). Mean modelled trophic levels of exclusively carnivores were estimated to be 3.42±0.16 and 3.47±0.13 at site B4S03 and B6S02, respectively, and the modelled trophic levels of exclusively deposit feeders were estimated as 2.55±8.08×10^−2^ and 2.50±8.59×10^−2^.

### 3.3 Carbon flows

Modeled total carbon input, i.e., the deposition of detritus and filter- and suspension feeding, had an estimated value of 6.84×10^−2^ mmol C m^−2^ d^−1^at site B4S03 and of 9.15×10^−2^ mmol C m^−2^ d^−1^ at B6S02 (Table 5). This input was dominated by phytodetritus deposition with a contribution of 58.3% (B6S02) to 62.1% (B4S03) to total deposition, followed by refractory detritus deposition (B4S03: 26.2%, B6S02: 33.0%). Additionally, an external pool of swimming copepods as prey for several predators (i.e. ctenophores, antipatharians, hydrozoans, and other anthozoans) contributed 4.78×10^−5^ mmol C m^−2^ d^−1^ (B6S02) to 4.01×10^−2^ mmol C m^−2^ d^−1^ (B4S03) to the modelled benthic ecosystem.

**Table 5.** Deposition of detritus (phytodetritus, semi-labile detritus, refractory detritus) and respiration of bacteria, metazoan meiobenthos, fish, and different macro- and megabenthos faunal groups at sites B4S03 and B6S02. Data are presented as flux values (mmol C m^−2^ d^−1^) and as % contribution to total deposition and total benthic respiration.

|  | B4S03 | % | B6S02 | % |
| --- | --- | --- | --- | --- |
| Total deposition | $6.84 \times 10^{-2}$ | 100 | $9.15 \times 10^{-2}$ | 100 |
| Phytodetritus | $4.25 \times 10^{-2}$ | 62.1 | $5.33 \times 10^{-2}$ | 58.3 |
| Semi-labile detritus | $7.94 \times 10^{-3}$ | 11.6 | $7.95 \times 10^{-3}$ | 8.68 |
| Refractory detritus | $1.80 \times 10^{-2}$ | 26.2 | $3.02 \times 10^{-2}$ | 33.0 |
| Total respiration | $8.57 \times 10^{-2}$ | 100 | $7.86 \times 10^{-2}$ | 100 |
| Bacteria | $3.78 \times 10^{-2}$ | 44.1 | $3.78 \times 10^{-2}$ | 48.1 |
| Metazoan meiobenthos | $1.04 \times 10^{-2}$ | 12.2 | $2.00 \times 10^{-2}$ | 25.4 |
| Macrobenthic polychaetes | $8.87 \times 10^{-3}$ | 10.4 | $3.06 \times 10^{-3}$ | 3.90 |
| Macrobenthic isopods | $3.31 \times 10^{-3}$ | 3.86 | $4.89 \times 10^{-3}$ | 6.23 |
| Macrobenthos (except polychaetes, isopods) | $9.68 \times 10^{-3}$ | 11.3 | $5.37 \times 10^{-3}$ | 6.84 |
| Megabenthic crustaceans | $1.38 \times 10^{-3}$ | 1.61 | $8.03 \times 10^{-4}$ | 1.02 |
| Megabenthic cnidarians | $4.65 \times 10^{-4}$ | 0.54 | $2.05 \times 10^{-3}$ | 2.60 |
| Holothurians | $3.99 \times 10^{-5}$ | $4.66 \times 10^{-2}$ | $4.27 \times 10^{-5}$ | $5.43 \times 10^{-2}$ |
| Poriferans | $8.10 \times 10^{-5}$ | $9.45 \times 10^{-2}$ | $8.89 \times 10^{-5}$ | 0.11 |
| Invertebrate megabenthos (except crustaceans, cnidarians, holothurians, poriferans) | $1.28 \times 10^{-2}$ | 15.0 | $3.60 \times 10^{-3}$ | 4.59 |
| Fish | $8.04 \times 10^{-4}$ | 0.94 | $9.18 \times 10^{-4}$ | 1.17 |

Most carbon was estimated to be lost from the two sampling sites via respiration (78.9–85.8% of total carbon loss), whereas carbon burial accounted for 1.30×10^−2^ mmol C m^−2^ d^−1^, and external predators predated with a rate of 9.89×10^−3^ mmol C m^−2^ d^−1^ at B4S03.

Estimated respiration ranged from 7.86×10^−2^ mmol C m^−2^ d^−1^ (B6S02) to 8.57×10^−2^ mmol C m^−2^ d^−1^ (B4S03). It was dominated by bacterial respiration that contributed between 44.1% (B4S03) and 48.1% (B6S02) to total respiration. Faunal respiration was estimated to be lower at B6S02 (4.08×10^−2^ mmol C m^−2^ d^−1^) than at B4S03 (4.79×10^−2^ mmol C m^−2^ d^−1^) and was dominated by metazoan meiobenthos and macrobenthos (except polychaetes and isopods) at B6S02 and by invertebrate megabenthos (except crustaceans, cnidarians, holothurians, and poriferans) and metazoan meiobenthos at B4S03 (Table 5). Modeled carbon ingestion of fauna is summarized in Supplementary figures 1 to 4. Estimated uptake of carbon by metazoan meiobenthos (Supplementary Fig. 1) was largest and accounted for 8.48×10^−2^ mmol C m^−2^ d^−1^ (B4S03) and 8.93×10^−2^ mmol C m^−2^ d^−1^ (B6S02).

Modeled carbon uptake by macrobenthos (Supplementary Fig. 2; B4S03: 4.08×10^−2^ mmol C m^−2^ d^−1^, B6S02: 3.84×10^−2^ mmol C m^−2^ d^−1^) and megabenthos (Supplementary Fig. 3; B4S03: 5.17×10^−2^ mmol C m^−2^ d^−1^, B6S02: 3.46×10^−2^ mmol C m^−2^ d^−1^) were of similar magnitude. The estimated carbon uptake of fish (Supplementary Fig. 4) was, however, one order of magnitude lower (B4S03: 2.69×10^−3^ mmol C m^−2^ d^−1^, B6S02: 1.72×10^−3^ mmol C m^−2^ d^−1^).

6.26–8.68% of total faunal ingestion was macrobenthic polychaetes’ carbon uptake, 4.00–7.79% macrobenthic isopods’ carbon uptake, and 9.36–9.59% all other macrobenthos’ carbon uptake. Estimated megabenthic carbon uptake was mediated for 1.28–4.02% of total faunal ingestion by megabenthic crustaceans, for 0.78–4.63% by megabenthic cnidarians, for 3.77×10^−2^–8.70% by holothurians, by 0.24–1.12% by poriferans, and for 5.38–23.1% by all other invertebrate megabenthos.

Uptake of DOC by bacteria as part of the microbial loop (see section 3.4) was 0.38 mmol C m^− 2^ d^−1^.

### 3.4 Carbon cycling and specific C pathways

*T*‥ was estimated to be 1.24 mmol C m^−2^ d^−1^ at site B4S03 and 1.20 mmol C m^−2^ d^−1^ at B6S02 and *FCI* was modeled to range from 0.51 mmol C m^−2^ d^−1^ (B4S03) to 0.53 mmol C m^−2^ d^−1^ (B6S02). The microbial loop, i.e., dissolution of detritus, uptake of DOC by bacteria, virus-induced bacterial mortality, bacterial respiration, and faunal grazing upon bacteria, had an estimated carbon flow of 0.84 mmol C m^−2^ d^−1^, which was 67.7% of modelled *T*‥ at B4S03 and 69.8% of modelled *T*‥ at B6S02.

An estimated carbon flow of 4.46×10^−3^ mmol C m^−2^ d^−1^ was channeled through the scavenging loop (i.e. scavenging of scavengers on carrion) at B4S03, which was 0.36% of modelled *T*… At B6S02, the carbon flow through the scavenging loop was modeled to be 3.32×10^−3^ mmol C m^− 2^ d^−1^ (0.28% of modelled *T*‥).

### 3.5 Consequences of polymetallic nodule removal

Excluding fauna from the food webs that have been predicted to be lost when polymetallic nodules are removed (Stratmann et al., 2021), led to a decrease in number of food-web compartments of 5.19% (B4S03) to 6.94% (B6S02) and to a loss of 4.61% (B6S02) to 8.88% (B4S03) of all links. Link density was reduced by 3.88% at B4S03 and increased by 2.51 at B6S02, and the connectance increased by 0.37% (B4S03) to 11.2% (B6S02). *T*‥ decreased by 2.85×10^−2^% at B4S03 and increased by 6.93×10^−2^% at B6S02, and *FCI* decreased by 0.34% (B4S03) and increased by 0.18% (B6S02), respectively. Network indices calculated for both sites in the presence and absence of nodule-related fauna are reported in Table 4.

A summary of the most important changes in modelled carbon flows (ΔC_flow_) due to the exclusion of nodule-dependent fauna is visualized in Fig. 5. At site B4S03, the largest changes in carbon flows were related to increased external predation upon invertebrate megabenthos (+8.21×10^−3^ mmol C m^−2^ d^−1^) and decreased scavenging activity of macrobenthos (−3.14×10^−3^ mmol C m^−2^ d^−1^). In comparison, at B6S02, the largest change in carbon flows was related to the increased sedimentary detritus uptake by metazoan meiobenthos (+6.29×10^−3^ mmol C m^−2^ d^−1^) and increased predation of megabenthic cnidarians on macrobenthic isopods (+3.51×10^−3^ mmol C m^−2^ d^−1^).

**Figure 5:**
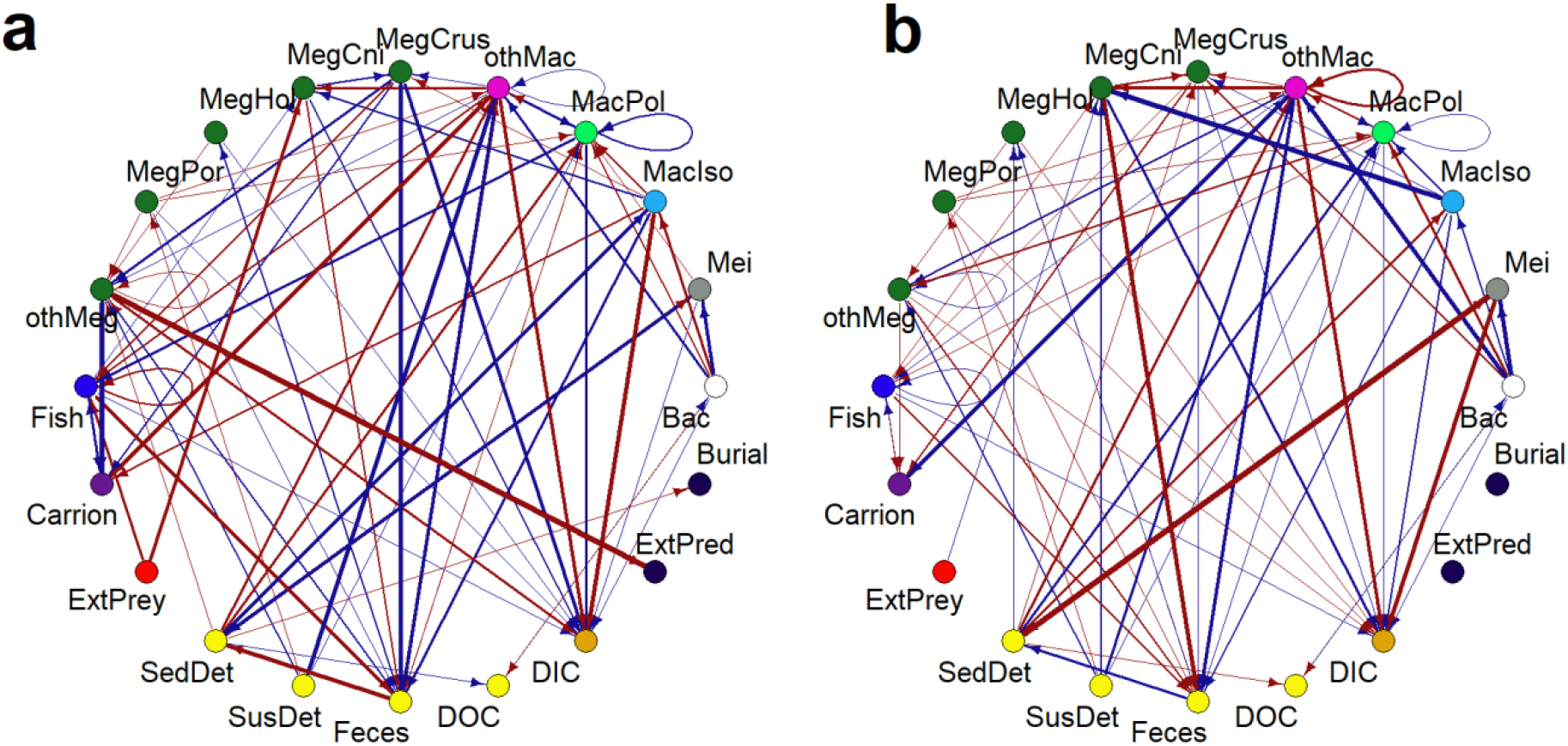
Figure summarizing changes in carbon flows between compartments as a result of the exclusion of polymetallic nodule-dependent fauna at sites a) B4S03 and b) B6S02. The width of the arrow corresponds to the double-square root of ΔC_flow_ (in mmol C m^−2^ d^−1^), where 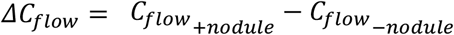, and the color indicates whether an exclusion of the fauna led to an increase in a specific carbon flow (red) or a decrease (blue). Abbreviations are: Bac = bacteria, ExtPred = external predation, ExtPrey = external prey, MacIso = macrobenthic isopods, MacPol = macrobenthic polychaetes, MegCni = megabenthic cnidarians, MegCrus = megabenthic crustaceans, MegHol = holothurians, MegPor = poriferans, Mei = metazoan meiobenthos, othMac = other macrobenthos, othMeg = invertebrate megabenthos, SedDet = sedimentary detritus, SusDet = suspended detritus.

Removing the nodule-dependent fauna did not affect the strength of the microbial loop. In contrast, the scavenging loop at B4S03 (-nodule) (7.09×10^−4^ mmol C m^−2^ d^−1^) was only 15.9% of the original scavenging loop and at B6S02 (-nodule), the scavenging loop accounted for 45.4% (= 1.51×10^−3^ mmol C m^−2^ d^−1^) of the initial scavenging loop.

## 4. Discussion

The role of polymetallic nodule-dependent fauna on benthic carbon cycling in the eastern CCZ was assessed by investigating differences in modelled carbon flows of intact food webs and of food webs where nodule-dependent fauna was excluded in the models. The food webs for two sites in the GSR exploration license area contained between 11 mmol C m^−2^ and 67 mmol C m^−2^ faunal carbon and between 20 mmol C m^−2^ and 35 mmol C m^−2^ bacterial carbon in 62 to 67 biotic compartments (excluding detritus, DOC, and carrion) (Fig. 3, Fig. 4, Table 1). The food webs were estimated to be fueled to 58% to 62% by phytodetritus deposition and to 26% to 33% by refractory detritus deposition, whereas most of the carbon was lost from the food webs as respiration. This respiration was dominated by bacterial respiration, followed by metazoan meiobenthos, and other macrobenthos respiration. Removing nodule-dependent faunal compartments had almost no effect on total system throughput *T*‥, but reduced the total number of links by 5% to 9% (Table 4).

### 4.1 Model limitations

Like all scientific models, also these highly resolved food-web models have limitations. Biomasses of metazoan meiobenthos, macrobenthos, invertebrate megabenthos, and fish were estimated by converting densities reported by the authors of the original peer-reviewed studies (see section 2.4.1) to carbon stocks using conversion factors (Supplementary tables 1 to 3). de Smet et al. (2021) assessed invertebrate megabenthos community composition and densities using autonomous underwater vehicle imagery obtained from ∼6 m above the seafloor and only reported specimens >5 cm. In a method paper, Schoening et al. (2020) compared abyssal faunal densities on seafloor images taken at 1.6 to 1.7 m above the seafloor with images taken 4.5 and 7.5 m above the seafloor and found three to 28 times higher densities in the low altitude images compared to the high altitude images. Consequently, in this study invertebrate megabenthos biomasses and subsequently their role in carbon cycling are underestimated.

Polymetallic nodules in abyssal plains have microbial communities inside their crevices (Blöthe et al., 2015; Cho et al., 2018) and on their surfaces (Wang et al., 2009) that are different from the microbial community in adjacent sediments (Cho et al., 2018; Molari et al., 2020). Hence, for a proper assessment of the role of polymetallic nodules on abyssal carbon cycling, not only the impact of nodule-dependent fauna exclusion, but also the effect of nodule-dependent microorganism exclusion on carbon cycling should be assessed. However, so far, no study has quantified the role of these microorganisms in carbon cycling and data on bacterial biomass inside and outside the nodules are lacking for the GSR exploration license area or any other license area in the CCZ. Therefore, I could not confidently include this process in the food web models for the sites B4S03 and B6S02, and used one (sedimentary) bacterial carbon pool per site instead.

Sweetman et al. (2019) measured 0.10±0.03 mmol C m^−2^ d^−1^ DIC uptake by benthic bacteria in the UK Seabed Resources Ltd (UK1) exploration license area and the Ocean Minerals of Singapore exploration area in the eastern CCZ. This value was 1.2 to 2.0 times higher than the uptake of phytodetritus carbon by bacteria at the sites (Sweetman et al., 2019) and therefore likely a very substantial contribution to abyssal carbon cycling. However, as the researchers conducting the baseline studies for GSR at B4S03 and B6S02 did not investigate bacterial dissolved inorganic carbon uptake, I am hesitant to include this process in the model (see also discussion in de Jonge et al. (2020)). Further studies in dark carbon fixation at different sites of the CCZ would therefore provide valuable information about the underlying mechanisms and which microorganisms are involved in them, and allow to include this process in future abyssal carbon-based food-web models.

### 4.2 Regional differences in carbon cycling at abyssal plains

The presented food webs estimated a difference in total system throughput *T*‥ of 3.75×10^− 2^ mmol C m^−2^ d^−1^ between sites B4S03 and B6S02, whereupon more carbon was recycled at B6S02 (Finn’s cycling index *FCI*) compared to B4S03 (Table 4). This difference in *T*‥ was partly the result of higher carbon cycling by deposit-feeding metazoan meiobenthos at B4S03 compared to B6S02 (Table 1) and might be related to small-scale variability in environmental parameters between the two sites. Metazoan meiobenthos densities in surface sediments (0–5 cm sediment depth) of the CCZ are influenced by water depth, silt content, and polymetallic nodule density (Hauquier et al., 2019). In a comparison of metazoan meiobenthos across various locations in the eastern CCZ, Hauquier et al. (2019) measured the highest meiobenthos density at a location with low nodule density, high pigment concentrations as proxy for (fresh) phytodetritus availability, and low clay content compared to the other sampling locations. Further sediment characteristics that influence meiobenthos distribution and densities in the eastern CCZ were identified in a modeling exercise by Uhlenkott et al. (2021). They include total organic and inorganic carbon content of the sediment, dry bulk sediment density, and shear strength. In this study, site B4S03, where higher metazoan biomass was detected, was indeed characterized by decreased nodule abundance compared to the B6S02 site, though mud content was higher and total organic carbon content was lower at B4S03 than at B6S02 (de Smet et al., 2017). Hence, I conclude that one important factor affecting carbon cycling on regional scale (280 km) in the eastern CCZ is environmental parameters.

Another reason for the differences in *T*‥ was the presence of Ctenophora at B4S03 whose biomass contributed 96% to total invertebrate megabenthic biomass at B4S03; they were not detected at B6S02 (Table 1). Ctenophora, also known as comb jellies, have been observed swimming in the UK Seabed Resources Ltd (UK1) exploration license area in the eastern CCZ (Amon et al., 2017), in the area of particular environmental interest (APEI) 6 in the northern CCZ (Jones et al., 2021; Simon-Lledó et al., 2019a), and in the exclusive economic zone (EEZ) of Kiribati in the western CCZ (Simon-Lledó et al., 2019b). They are carnivorous (Haddock, 2007) and predate upon krill (Swift et al., 2009) and copepods (Toyokawa et al., 2003). Therefore, I included an external pool of abyssopelagic copepods as prey for Ctenophora in the food-web model for site B4S03. This model estimated that Ctenophora ingested 3.95×10^−2^ mmol C m^−2^ d^−1^ abyssopelagic copepods and 9.31×10^−4^ mmol C m^−2^ d^−1^ macrobenthic copepods which is equivalent to 3.27% of *T*‥ at B4S03. In contrast to the difference in metazoan meiobenthos biomass between the two sampling sites at the GSR exploration license area, which was likely caused by variation in environmental parameters, I expect that the absence of mobile Ctenophora at B6S02 was related to their low density in the CCZ (<1.30×10^−3^ ind. m^−2^ at Belgian license area (de Smet et al., 2021), <0.03 ind. m^−2^ at APEI6 (Simon-Lledó et al., 2019a)) and not to any particular environmental (or other) parameter.

### 4.3 Assessment of polymetallic nodule removal on C cycling and outlook to prospective deep-seabed mining effects

Excluding nodule-dependent fauna from the food webs developed for sites B4S03 and B6S02 impacted the faunal contribution to *T*‥, but had no effect on the microbial loop. The taxa, that were removed, belonged to filter and suspension feeders taking up marine snow (Brachiopoda, Crinoidea), but also to scavengers (Amphipoda), and carnivorous suspension feeders (Brisingida, Actinaria) predating upon small crustaceans (Table 1, Supplementary table 2, Supplementary table 3).

Scavenging amphipods were responsible for 66% of scavenging occurring at B4S03, so that their absence as a result of nodule removal strongly reduced the scavenging loop. Scavengers in the CCZ include demersal fish, such as *Coryphaenoides* spp., *Bassozetus* sp., and *Pachycara nazca*, and invertebrate megabenthos, like the amphipod *Eurythenes gryllus*, squat lobster *Munidopsis* sp., and the shrimp *Cerataspis monstrosus* (Harbour et al., 2020). Their taxon density and diversity vary regionally and Drazen et al. (2021) observed that the scavenger assemblage in the eastern CCZ was very different from the assemblage in the western APEIs due to higher abundances of the shrimp *Hymenopenaeus nereus* and lower abundances of the fishes *Barathrites iris* and *Coryphaenoides* spp‥ The latter fish was also observed in the GSR exploration license area (de Smet et al., 2021) and the carbon-based food-web model for B4S03 estimated that *Coryphaenoides* spp. partly compensated the loss of scavenging amphipods by increasing its contribution to the scavenging loop from 23.5% to 56.6%.

Abyssal scavengers react relatively fast to the deposition of carrion at the seafloor: *Coryphaenoides* spp. were attracted to fish bait (i.e. albacore tuna) at the seafloor of the eastern CCZ within 1 h (Harbour et al., 2020)) and synaphobranchid eels reached the fish bait (i.e. Pacific mackerel) in the western CCZ after ∼7 h (Leitner et al., 2021, 2020). Also during a natural mass deposition event of gelatinous pyrosome carcasses in the Peru Basin in 2015, actinarians, ophiuroids, asteroids, and isopods were observed to (putatively) scavenge on the pyrosomes carcasses within a couple of days (personal observation, Hoving et al. (in review)). Hence, an industrial deep-seabed mining operation will likely attract scavenging fish and invertebrate megabenthos within hours. This will overcompensate the loss of nodule-dependent scavengers and the importance of the scavenging loop for benthic carbon cycling will strongly increase, unless scavengers are chased off by the sediment plume or by the toxic metals that might be released during the mining operation (Hauton et al., 2017; Koschinsky et al., 2001; Miller et al., 2018).

At B4S03, filter and suspension feeding Copepoda and Bivalvia benefitted from the loss of nodule-dependent filter and suspension feeding Brachiopoda and Crinoidea by ingesting 28 to 48% more labile marine snow. In comparison, more labile marine snow particles, that nodule-dependent filter and suspension feeders normally would have filtered out of the water column, settled on the sediment at B6S02 where they contributed to the sedimentary phytodetritus pool. As a result, epistrate feeding, selective and non-selective deposit-feeding nematodes were estimated to consume more labile phytodetritus at this site, though the contribution of phytodetritus to their overall diet changed only minimally.

As the presence of polymetallic nodules reduces the substrate availability for metazoan meiobenthos not dependent on polymetallic nodules, nodule-free areas in the GSR exploration license area have a higher nematode density than nodule-rich areas (Pape et al., 2021). Deep-sea metazoan meiobenthos occurs predominantly in the upper 5 cm of sediment, whereupon >60% live in the surface 1 cm and <10 % in the 4–5cm layer (Vincx et al., 1994). During an industrial deep-seabed mining operation, however, the upper 4–8 cm of surface sediment will be removed (Haeckel, 2022). Consequently, meiobenthos will likewise be removed and cannot benefit from the potentially increased access to labile detritus, resulting in a loss of 11% (= 0.13 mmol C m^−2^ d^−1^) of benthic carbon cycling at B4S03 and 12% (= 0.14 mmol C m^−2^ d^−1^) of benthic carbon cycling at B6S02.

The top 1 m of seafloor sediment is assumed to contain 10^8^ prokaryotic cells cm^−3^ (Jørgensen and Boetius, 2007) and Nomaki et al. (2021) measured prokaryotic cell abundances of 3.6×10^7^ cells g^−1^ in the 0–1 cm depth layer to 1.4×10^7^ cells g^−1^ in the 5–10 cm depth layer in the abyssal west Pacific. In the eastern CCZ, bacterial biomasses range from 238 mg C m^−2^ to 530 mg C m^−2^ (Pape et al., 2017; Sweetman et al., 2019) in the upper 5 cm of sediment which will be removed during industrial deep-seabed mining operations. As the models in this study calculate, the microbial loop in the upper 5 cm of sediment contributes between 68% and 70% to *T*‥ at the GSR exploration license area. Hence, we can expect that an impairment of said microbial loop during deep-seabed mining operations will have the largest impact on benthic carbon cycling. Vonnahme et al., (2020) measured an up to fourfold lower prokaryotic activity in a 5 week old epibenthic sledge track in the more eutrophic (compared to the CCZ) abyssal Peru Basin. When I assume that mining-induced sediment removal will have a similar, if not stronger effect on the microbial loop in the CCZ, benthic carbon cycling at the GSR exploration license area will be further reduced by 0.63 mmol C m^−2^ d^−1^. Hence, this study indicates that deep-seabed mining in the GSR license area will likely reduce ecosystem function in the form of carbon cycling by minimum 61 to 64%. Further expected loss of nodule-independent macrobenthos and other invertebrate megabenthos has not been taken into account.

## 5. Conclusion

Based on very highly-resolved abyssal food-web models for two sites in the GSR exploration license area, I show that benthic carbon cycling in the eastern CCZ varies regionally (280 km) in the same order of magnitude as benthic carbon cycling 26 years after a small-scale sediment disturbance experiment between reference sites and areas likely affected by re-sedimentation (i.e., sites called ‘outside the plough tracks’ in de Jonge et al. (2020)). Hence, the International Seabed Authority should request a spatially (and temporally) very homogenous and fine-scale (environmental baseline) sample coverage for each sub-exploration license area in the CCZ to be able to assess serious harm to the marine environment (Levin et al., 2016)and to determine potential recovery from prospective deep-seabed mining later on. The models furthermore indicate that polymetallic nodule-dependent fauna has a very minor contribution to benthic carbon cycling, though it is important for marine biodiversity (Niner et al., 2018; Stratmann et al., 2021) and therefore the author recommends to concentrate on the microbial loop when investigating whether this specific ecosystem function is affected by deep-seabed mining, and on marine biodiversity.

## Declaration of Competing Interest

The author declares that she has no competing financial interest or personal relationship that could have appeared to influence the work reported in this paper.

## Acknowledgements

This study received funding by JPI Oceans – Impacts of deep-sea nodule mining project “Mining Impact 2” from the Dutch Research council (NWO-ALW grant 856.18.003), from the research program NWO-Rubicon with project number 019.182EN.012, and from the NWO-Talent program Veni with project number VI.Veni.212.211.

## Appendix A

This appendix contains further information and data used to construct the food-web inverse models.

**Table A.1:** Summary of size-class specific faunal taxa (total number of lost taxa, % of phyla lost) from the Clarion-Clipperton Fracture Zone that are estimated to be lost due to the removal of polymetallic nodules (data from Stratmann et al. (2021)).

| Size class | Phyla | Class | Order |
| --- | --- | --- | --- |
| Meiobenthos | Arthropoda (1, 1.18%) |  |  |
|  | Cnidaria (1, 50.0%) |  |  |
|  | Foraminifera (6, 6.82%) |  |  |
| Macrobenthos | Annelida (1, 0.76%) | Polychaeta (1, 0.77%) |  |
|  | Arthropoda (16, 12.6%) | Malacostraca (17, 14.8%) | Amphipoda (13, 100%)* |
|  |  |  | Isopoda (1, 1.67%) |
|  |  |  | Tanaidacea (2, 5.88%) |
|  | Brachiopoda (2, 66.7%)* | Rhynchonellata (2, 50.0%) |  |
|  |  | Lingulata (1, 100%) |  |
|  | Bryozoa (17, 81.0%) | Gymnolaemata (7, 87.5%) |  |
|  |  | Stenolaemata (10, 76.9%) |  |
|  | Chordata (1, 100%) | Ascidacea (1, 100%) |  |
|  | Cnidaria (4, 57.1%) | Anthozoa (2, 40.0%) |  |
|  |  | Hydrozoa (1, 100%) |  |
|  |  | Scyphozoa (1, 100%) |  |
|  | Echinodermata (1, 20.0%) | Crinoidea (1, 100%) |  |
|  | Foraminifera (43, 91.5%) |  |  |
|  | Mollusca (3, 11.1%) | Bivalvia (1, 6.67%) |  |
|  |  | Gastropoda (1, 25.0%) |  |
|  | Platyhelminthes (1, 100%) | - (1, 100%) |  |
|  | Porifera (1, 50.0%) | Demospongiae (1, 100%) |  |
| Invertebrate Megabenthos | Annelida (1, 9.09%) | Polychaeta (1, 9.09%) |  |
|  | Arthropoda (6, 25.0%) | Malacostraca (5, 23.8%) | Amphipoda (3, 60.0%) |
|  |  |  | Isopoda (1, 50.0%) |
|  |  |  | Mysida (1, 100%) |
|  |  | Thecostraca (1, 100%) |  |
|  | Chordata (1, 11.1%) | Ascidacea (1, 12.5%) |  |
|  | Cnidaria (11, 25.6%) | Anthozoa (8, 27.6%) | Actiniaria (11, 100%)* |
|  | Echinodermata (15, 16.7%) | Asteroidea (2, 12.5%) | Brisingida (2, 66.7%)* |
|  |  | Crinoidea (3, 75.0%)* |  |
|  |  | Ophiuroidea (10, 41.7%) |  |
|  | Porifera (1, 3.33%) | Hexactinellida (1, 4.55%) | Lyssacinosida (1, 6.25%) |
| Fish | Chordata (1, 6.25%) | Actinopteri (1, 100%) | Perciformes (1, 100%) |

\*Taxa (on phyla, class, or order level) that were observed in the GSR exploration license area and are part of the present food-web model. These taxa were excluded in the models developed for section 3.5 when $\geq 60\%$ of the specific phyla/ class/ order would be lost if nodules were removed.

**Table A.2:** List of food-web compartments for which specific physiological process constraints [assimilation efficiency *AE* (-), net growth efficiency *NGE* (-), secondary production *SP* (mmol C m^−2^ d^−1^), respiration *R* (mmol C m^−2^ d^−1^), and feeding preferences *FP* (-)] were adjusted in the fitting step. The abbreviations of the food-web compartments are listed in Table 1.

| Physiological process | Adjusted constraint | Food-web compartments at B4S03 (+ nodules) | Food-web compartments at B4S03 (- nodules) | Food-web compartments at B6S02 (+ nodules) | Food-web compartments at B6S02 (- nodules) |
| --- | --- | --- | --- | --- | --- |
| *AE* | [0.18, 1.00] |  |  | MeiDF | MeiDF |
|  | [0.10, 0.89] | Mac_Oli, Pol_Eup, Mac_otherCru, Mac_Biv | Mac_Oli, Pol_Eup, Mac_otherCru, Mac_Biv |  |  |
|  | [0.68, 1.00] | Pol_Gly, Pol_Goni, Pol_Lumb, Mac_Pyc | Pol_Gly, Pol_Goni, Pol_Lumb, Mac_Pyc |  |  |
|  | [0.10, 0.75] |  |  | Meg_Ant, Meg_othAs, Meg_Cri, Meg_Asp, Meg_Elp, Meg_othEl, Meg_unHo, Meg_Cho, Meg_Doc, Meg_Eup, Meg_othPor | Meg_othAs, Meg_Asp, Meg_Elp, Meg_othEl, Meg_unHo, Meg_Cho, Meg_Doc, Meg_Eup, Meg_othPor |
|  | [0.40, 1.00] |  |  | Meg_Oph | Meg_Oph |
|  | [0.10, 1.00] |  |  | Meg_Alc, Meg_Cer, Meg_othAn, Meg_Hyd, Meg_Bri | Meg_Alc, Meg_Cer, Meg_othAn, Meg_Hyd |
|  | [0.10, 0.87] | Fish_Cor | Fish_Cor | Fish_Ost | Meg_Ant, Fish_Ost |
| *NGE* | [0.10, 1.00] |  |  | MeiDF |  |
|  | [0.10, 0.68] | Pol_Goni, Pol_Mal,<br>Pol_Nep, Pol_Ner,<br>Pol_Ophe,<br>Pol_Siga, Pol_Syl,<br>Pol_Tri,<br>Mac_Cope,<br>Mac_Amp, IsoDF,<br>IsoNoSDF,<br>IsoSeDF,<br>Mac_Mys,<br>Mac_Ost,<br>Mac_Biv,<br>Mac_Cha,<br>Mac_Nema,<br>Mac_Neme | Pol_Goni, Pol_Mal,<br>Pol_Nep, Pol_Ner,<br>Pol_Ophe,<br>Pol_Siga, Pol_Syl,<br>Pol_Tri,<br>Mac_Cope, IsoDF,<br>IsoNoSDF,<br>IsoSeDF,<br>Mac_Mys,<br>Mac_Ost,<br>Mac_Biv,<br>Mac_Cha,<br>Mac_Nema,<br>Mac_Neme |  |  |
|  | [0.10, 0.87] |  | Fish_Cor |  |  |
|  | [0.23, 1.00] |  |  | Meg_Oph,<br>Meg_Cho,<br>Meg_Doc,<br>Meg_Eup,<br>Meg_othPor | Meg_unHo,<br>Meg_Oph,<br>Meg_Cho,<br>Meg_Doc,<br>Meg_Eup,<br>Meg_othPor |
| *SP* | $[2.57 \times 10^{-3}, \infty] \times C$<br>stock | | | Pol_Lumb | |
| | $[1.00 \times 10^{-5}, 1.47 \times 10^{-3}] \times C$<br>stock | Meg_Ant | Meg_Ant | | |
| *R* | $[1.26 \times 10^{-3}, \infty] \times C$<br>stock | Meg_Act,<br>Meg_Ant | Meg_Ant | | |
| | $[1.00 \times 10^{-8}, 1.26 \times 10^{-3}] \times C$<br>stock | | | Meg_othAn | Meg_othAn |
| | $[0, 1.26 \times 10^{-3}] \times C$<br>stock | | | | |
| *FP* | [0.10, maxfracPupt] | Pol_Syl, Meg_Act |  | Meg_Oph | Meg_Oph |
|  | [0.25, maxfracPupt] | Pol_Phy, Mac_Pyc,<br>Meg_Gal,<br>Meg_othDec,<br>Meg_Asp,<br>Meg_othCni | Pol_Phy, Mac_Pyc |  |  |
|  | [0.01, maxfracPupt] | Mac_Tan,<br>Mac_Oph,<br>Meg_Oph | Mac_Tan,<br>Mac_Oph,<br>Meg_Oph | Mac_Tan | Mac_Tan |

**Table A.3:** Fish-taxon specific dietary constraints [min, max] for the two sampling sites B4S03 and B6S02. The contribution of carrion to the fish diet was not specifically constrained.

| **Fish taxon** | **Prey** | **B4S03** | **B6S02** |
| --- | --- | --- | --- |
| *Coryphaenoides* spp. | *Ipnops* sp. | [0.00, 1.00] | [0.00, 0.05] |
|  | Aristeidae | [0.10, 0.15] | [0.00, 0.03] |
|  | Galatheidae | [0.10, 0.53] | [0.40, 0.50] |
|  | Other megabenthic Decapoda | [0.015, 0.04] | [0.00, 0.01] |
|  | Macrobenthic Polychaeta | [0.35, 0.40] | [0.10, 0.16] |
| *Ipnops* sp. | Crustaceans | [0.25, 0.35] | [0.25, 1.00] |
|  | Aristeidae | [0.10, 0.14] | [0.00, 0.04] |
|  | Galatheidae | [0.40, 0.51] | [0.50, 0.60] |
|  | Other megabenthic Decapoda | [0.00, 0.03] | [0.00, 0.01] |
| Osteichthyes morphotypes | *Ipnops* sp. | [0.025, 0.07] | [0.025, 0.06] |
|  | Aristeidae | [0.04, 0.08] | [0.00, 0.04] |
|  | Galatheidae | [0.20, 0.29] | [0.45, 0.56] |
|  | Other megabenthic Decapoda | [0.00, 0.10]* | [0.00, 0.10]* |
|  | *Coryphaenoides* spp. | [0.45, 0.54] | [0.25, 0.33] |

\*Constraint adjusted in the fitting step.

## Appendix B. Supplementary material

Supplementary data to this article can be found online at XXX.

## Notes

### Competing Interest Statement

The authors have declared no competing interest.

